# Inducing TRIB2 Targeted Protein Degradation to Reverse Chemoresistance in Acute Myeloid Leukaemia

**DOI:** 10.1101/2025.10.03.680259

**Authors:** Evie Rigby, Akshara Narayanan, Elzbieta Kania, John A Harris, Jamie Williams, Binghua Zhang, Lijun Liu, Laura Richmond, Fengtao Zhou, Ke Ding, Ruaidhrí J Carmody, Patrick A Eyers, Karen Keeshan

**Affiliations:** Wolfson Wohl Translational Cancer Research Centre, School of Cancer Science, College of Medical, Veterinary and Life Sciences, University of Glasgow, UK; Department of Biochemistry, Cell and Systems Biology, Institute of Systems, Molecular and Integrative Biology, University of Liverpool, UK; School of Molecular Biosciences, College of Medical, Veterinary & Life Sciences, Davidson Building, Room, University of Glasgow, UK; International Cooperative Laboratory of Traditional Chinese Medicine Modernization and Innovative Drug Development, Ministry of Education of People’s Republic of China, College of Pharmacy, Jinan University, China; Shanghai Institute of Organic Chemistry, Chinese Academy of Sciences, China

## Abstract

The myeloid oncogene TRIB2 is a key driver of acute myeloid leukaemia (AML) pathogenesis, promoting chemoresistance and blocking differentiation through ubiquitin-mediated degradation of the C/EBPα transcription factor. Despite its stable and sometimes elevated expression across AML subtypes, TRIB2 remains a clinically-untargeted vulnerability. Here, we present a comprehensive investigation into TRIB2 degradation mechanisms using multimodal approaches, including CRISPR knockout, mutational protein stability, small molecule TRIB2 engagement and evaluation of a novel targeted protein degrader (TRIB2-PROTAC). We identify Afatinib, a multi-ERBB covalent inhibitor, as a rapid inducer of TRIB2 degradation, triggering AML cell death via an ERBB-independent pathway. Importantly, TRIB2 degradation synergized with cytarabine, the frontline AML chemotherapy, amplifying therapeutic efficacy. Mapping of TRIB2 ubiquitination sites revealed Lys-63 as critical for its own proteolytic turnover, and a Lys to Arg degradation-resistant mutant (K*all*R) conferred enhanced chemoresistance and increased leukaemic engraftment *in vivo*. CRISPR-mediated TRIB2 knockout validated an essential role in AML cell survival. Consistently, the novel TRIB2-PROTAC (compound 5K) achieved robust TRIB2 degradation and AML cell killing at low micromolar concentrations. These findings establish TRIB2 as a compelling therapeutic target in AML and demonstrate that leveraging the ubiquitin-proteasome system to degrade TRIB2 offers a promising strategy to overcome chemoresistance. This work provides strong preclinical rationale for the development of TRIB2-targeting therapies in AML.

## Introduction

Acute myeloid leukaemia (AML) represents one of the most aggressive haematological malignancies, characterised by the rapid proliferation of abnormal myeloid cells and poor long-term survival rates despite patient-intensive therapeutic intervention. The molecular complexity underlying AML pathogenesis has driven extensive research into novel therapeutic targets that can overcome the limitations of conventional chemotherapy and address the challenge of therapy resistance. While overall patient survival has improved in recent years, due to a growing understanding of disease pathogenesis and subsequent improvements to standards of supportive care, AML cases continue to rise. Relapse and treatment resistance is still a major unmet clinical need and there is an urgent need to identify novel targets and treatment strategies in AML.

Among emerging targets of interest, Tribbles pseudokinase 2 (TRIB2) has gained significant attention as an oncogene with well-established roles in leukaemogenesis. TRIB2 belongs to the Ser/Thr (pseudo)kinase family of proteins, regulating pleiotropic cellular processes through phosphorylation-independent mechanisms (1). Unlike canonical kinases, which phosphorylate substrates, TRIB2 functions as a conformationally- switchable protein adaptor, serving as an essential link in the ubiquitin-proteasome degradation system by facilitating interactions between target ‘substrates’ that are ubiquitinated by bound E3 ligases, increasing Lys- 48 levels of ubiquitinated proteins for proteasomal degradation in cells (2, 3). Beyond its role in protein homeostasis, TRIB2 contributes to cancer progression and therapy resistance through multiple adaptor functions. Indeed, TRIB2 has recently been implicated in regulating ferroptosis, an iron-dependent form of programmed cell death, that represents a potential therapeutic vulnerability in cancer cells. By facilitating βTrCP-mediated ubiquitination of the transferrin receptor (TFRC), TRIB2 reduces intracellular iron and desensitises cells to ferroptosis (4).

The oncogenic role of TRIB2 in AML has been firmly established through extensive mechanistic studies. TRIB2 functions as an AML-driving oncogene through its interaction with the E3 ubiquitin ligase COP1, which mediates the degradation of the transcription factor C/EBPα (2, 5, 6). This degradation pathway is particularly critical in AML pathogenesis, as C/EBPα is essential for normal myeloid differentiation. By promoting C/EBPα degradation, TRIB2 effectively blocks myeloid differentiation, leading to the accumulation of immature myeloid precursors characteristic of AML.

Our previous work provided compelling evidence that targeting TRIB2 depletion in AML represents a promising therapeutic strategy. Genetic knockdown of TRIB2 using shRNA-based approach in AML cells demonstrated significant therapeutic efficacy, leading to approximately 50% cell death and substantially impairing AML engraftment capabilities. Inhibition of proteasome-mediated degradation of C/EBPα reversed TRIB2 oncogenic function (7, 8). Furthermore, TRIB2 shRNA knockdown demonstrated synergistic effects when combined with standard AML chemotherapeutic agents suggesting that TRIB2 depletion could potentiate the efficacy of existing therapeutic regimens (9). Moreover, TRIB2 abundance has been linked to therapy resistance in several different cancer types including AML through the activation of pro-survival pathways and disruption of tumour suppressive pathways such as the BCL2 family, AKT-FOXO and MDM2/p53 pathways (9–12).

Building on the genetic validation of TRIB2 as a therapeutic target, efforts have focused on identifying pharmacological approaches for TRIB2 depletion. *In vitro* small molecule screening has revealed that multi-ERBB inhibitors possess the capacity to bind and degrade TRIB2 in cells (13). This discovery provided the first pharmacological proof-of-concept for TRIB2-targeted therapy in AML. The mechanism underlying this effect involves covalent interaction of second and third generation dual HER2 and EGFR inhibitors, including afatinib, with unique cysteine residue(s) present in regulatory regions of the TRIB2 pseudokinase domain. Importantly, interaction is specific for TRIB2 due to the absence of equivalent cysteine residues in related pseudokinases, including TRIB1, TRIB3, and STK40/SGK495.

The stability and regulation of TRIB2 present both challenges and opportunities for therapeutic intervention. TRIB2 stability is itself actively regulated through complex interactions with E3 ubiquitin ligases and the proteasome system. The SCFβ-TRCP ubiquitin ligase complex, composed of the F-box protein β-TRCP, scaffold protein CUL1, and adapter protein SKP1, polyubiquitinates TRIB2 at its N-terminal degradation domain, triggering proteasomal degradation (14). TRIB2 Lys residues are therefore predicted to be crucial for downstream E3 ligase-mediated ubiquitination events affecting target protein degradation and signalling outputs. Understanding the relevant sites of ubiquitination that control TRIB2 stability will provide key insights into potential strategies for modulating TRIB2 levels therapeutically. Interestingly, TRIB2 has also been shown to influence cellular ubiquitin levels by indirectly modulating Proteasome 20S Subunit Beta 5 (PSMB5) (15) thereby affecting protein homeostasis, a process increasingly recognized as a contributor to AML pathogenesis. Thus, TRIB2 serves as an increasingly-attractive target for modulation in this disease. This study aimed to evaluate the cellular efficacy and tolerability of targeted TRIB2 degradation in AML cells, with the goal of establishing preliminary clinical evidence for TRIB2-directed therapies in this aggressive disease.

## Materials and Methods

### Cell Lines and Culture

All cell lines were obtained commercially and cultured according to standard mammalian tissue culture protocols (9). AML suspension cell lines were grown at 37° C, 5% CO_2_ and maintained at appropriate cell densities in Rosewell Park Memorial Institute (RPMI)-1640 culture media (Invitrogen) containing 2mM L-Glutamine, 100 I.U/mL Penicillin, 100 µg/mL streptomycin and supplemented with relevant concentrations of foetal bovine serum (FBS). Human embryonic kidney (HEK)-293T cells were cultured at 37° C, 5% CO_2_ in appropriate Dulbecco Modified Eagle Medium (DMEM; Invitrogen) containing 2mM L-Glutamine, 100 I.U/mL Penicillin, 100 µg/mL streptomycin and 10% (v/v) FBS. Cell growth and viability was monitored via trypan blue counts. Cells were periodically screened for mycoplasma infection.

### Cell viability and apoptosis assays

Resazurin assays were employed to evaluate metabolic activity and cell viability. Cells were seeded in 96-well plates 24 hours prior to compound addition. Compounds solubilised in DMSO, including Afatinib and PROTAC compound 5K, were added to plates at relevant concentrations. Resazurin was added at a final concentration of 50µM and incubated for 4 hours. Absorbance was determined on a TECAN infinite M200 PRO plate reader. Cell viability was calculated through normalisation to untreated controls. For apoptosis assays to evaluate cell death, after defined treatments, cells were washed twice with ice cold PBS, then resuspended in binding buffer containing annexin V-phycoerythrin (BD Biosciences) and 4’6 diamidino-2- phenylindole (DAPI; Sigma-Aldrich). Samples were incubated for 15 minutes in the dark prior to flow cytometry analysis.

### Drug assays

AraC was resuspended in water. PROTAC 5K was synthesised (16) and resuspended in DMSO. Afatinib was obtained from LC laboratories (cat A-8644) and resuspended in DMSO. AML cell lines were treated concurrently with AraC and afatinib at various concentrations to determine synergism. Cells were seeded in 96-well plates at a concentration of 2.5x10^4^ cells/well, 24 hours prior to compound addition. Drugs were added at the indicated final concentrations prior to cell viability assays. The SynergyFinder application was used to calculate synergy scores for drug combinations at each concentration, in accordance with the Bliss synergy model. Dose response matrices and synergy maps were generated through SynergyFinder (17).

### Molecular Modelling

The structure of human full length human TRIB2 (UniProt: Q92519) containing amino acids 1-343 was predicted using AlphaFold 3 (18) and the top ranked output (pTM = 0.77) was modelled using the molecular visualization program UCSF ChimeraX (19).

### Immunoblotting

For protein lysates, cells were washed twice in ice cold PBS and lysed with RIPA buffer (50mM TRIS-HCl pH 8.0, 1% NP40, 1% sodium deoxycholate, 150mM sodium chloride, 1mM EDTA) supplemented with cOmplete™ EDTA-free Protease Inhibitor Cocktail (Roche, Cat# 04693132001) and PhosSTOP™ (Roche, Cat# 4906837001). For whole cell lysates, 5x10^5^ cells were collected in 20µl 2x Sodium Dodecyl Sulfate (SDS) sample buffer, containing β-mercaptoethanol (BME), heated to 95°C and snap frozen. Western blotting was carried out according to standard procedures. Protein concentrations in RIPA-lysed samples were determined using Bradford or BCA Assays. Samples were supplemented with SDS sample buffer and boiled before separation on 10% polyacrylamide gels. Proteins were transferred onto nitrocellulose membranes and blocked with 5% non-fat skimmed milk or BSA in TBS-T. All primary antibodies (total-ERK 1:1000 in 5% BSA [CellSignaling: cat 4695s], phospho-ERK 1:1000 in 5% BSA [CellSignaling: cat 4370s], TRIB2 1:1000 in 2% milk [proteintech: cat 15359-1-AP], GAPDH 1:2000 in 5% milk [CellSignaling: cat 2118], ANTI-FLAG 1:1000 in 5% milk [Merck: cat F1804], ANTI-HA 1:1000 in 5% milk [SantCruz: cat 7392]) were incubated at 4°C overnight followed by three 10-minute TBS-T washes and 1 hour incubation in secondary antibody (1:5000 anti-Rabbit or anti-mouse in 5% milk [Invitrogen: cat 31460/31430) at room temperature. Proteins were visualised and quantified using ECL on a Licor Odyessy Fc.

### Cloning, Transfection and Retroviral transduction

To permit ubiquitination assays, a 1068 bp fragment encoding the entire human Trib2 cDNA was synthesised alongside a C-terminal FLAG eptitope tag prior to the stop codon, and subcloned into pcDNA3.1 via digestion with HindIII/XbaI (GenScript). TRIB2 Lys (K)-mutants were synthesised by *de novo* gene synthesis and subcloned into pcDNA3.1 vectors. Plasmid DNA for cellular transfection was prepared using standard procedures. For stably transduced TRIB2-expressing THP1 cell lines, full-length human Trib2 with a C-terminal FLAG epitope was subcloned into pIG (puro IRES GFP) with XhoI/HpaI restriction enzymes (GenScript). 1068bp fragments encoding TRIB2 Cys96Ser/Cys104Ser-substitutions or a fragment in which all Lys (K) residues were substituted with Arg (R) (K*all*R) TRIB2 were also synthesised and subcloned into pIG. For TRIB2 ubiquitination assays TRIB2 WT and K-mutant plasmids were transfected into HEK293T cells using Turbofect transfection reagent (ThermoFisher) according to the manufacturer’s protocol. After transfection, cells were incubated for a further 48 hours before treatment with MG132 for 2 hours, prior to cell lysis and protein extraction. To generate stably transduced THP1 cell lines, retrovirus was produced by transient transfection of TRIB2 WT, C96/104S or KallR mutant constructs in pIG backbones alongside the packaging vectors pCGP and VSVG into HEK293T cells using Turbofect. After a 24hr hour incubation, retrovirus-containing supernatants were harvested via centrifugation and filtration and used immediately or stored at -80°C prior to analysis. Media was replaced and supernatant similarly harvested the following day (48hr post-transfection). To transduce THP1 and U937 cells and generate stably transduced lines, 1x10^6^ cells/well were seeded in 12-well plates and incubated for 24 hours. Plates were centrifuged and media removed before cells were resuspended in 400µl fresh media. 1ml viral supernatant aliquots were added to cells in the presence of 4µg/ml polybrene. Viral transduction was performed via spinoculation at 700xg for 90 minutes at room temperature followed by overnight incubation at 37°C. Spinoculation was repeated the following day using 48hr viral supernatant and cells incubated in plates for 6 hours before transferring to T25 flasks supplemented with 9ml media for overnight incubation. The following day cells were cultured in media containing 2µg/ml puromycin for 48 hours, followed by puromycin reduction to 1µg/ml for maintenance. Transduction efficiency was verified using flow cytometry to calculate the percentage of GFP +ve cells using a BD Verse flow cytometer and confirmed by qPCR after total mRNA isolation.

### Ubiquitination assays

Ubiquitination assays were performed on lysates generated from TRIB2 WT/K-mutant transfected HEK293T cells. Transiently transfected cells were treated with 10µM MG132 for 2 hours before washing in ice cold PBS. Cells were lysed in denaturing lysis buffer (1% (w/v) SDS, 50mM sodium HEPES, 100mM sodium chloride, EDTA-free complete protease inhibitor mix, 1mM N-ethylmaleimide), boiled at 95°C for 5 min, and sonicated to shear DNA. Whole cell lysates were diluted 1:10 in non-denaturing lysis buffer (50mM sodium HEPES, 100mM sodium chloride, 1% (v/v) triton X-100, 0.5% (w/v) sodium deoxycholate, 1mM N-ethylmaleimide) and the supernatant isolated by centrifugation. Protein concentrations were determined using a standard BCA assay protocol. For immunoprecipitation, Dynabeads magnetic beads (ThermoFisher: cat 10003D) were first washed in non-denaturing lysis buffer. 20µl Dynabeads were used with 1mg total protein in 1ml non-denaturing lysis buffer per reaction and 1µl anti-FLAG antibody or IgG control antibody added. Samples were incubated at 4°C with rotation overnight. Beads were washed with non-denaturing lysis buffer and target protein eluted in sample buffer (4% SDS, 8M Urea, 50µM DTT) and boiled before immunoblotting analysis after SDS-PAGE.

### TRIB2 Stability assays

To assess the stability of key TRIB2 K-mutants, HEK293T cells transfected with TRIB2 constructs were either treated with 10µM MG132 or 100µM Cycloheximide (CHX) for 3 or 6 hours before RIPA lysis as previously described. Protein expression and stability was assessed via western blot using anti-FLAG A, with protein degradation over time compared with untreated controls.

### CRISPR and siRNA

Three sgRNA oligos and their complementary reverse sequences were designed for both TRIB2 and TRIB1, using GenScript designed sgRNA sequences calculated to have high on-target efficiency. Forward and reverse strands were purchased as single stranded DNA (ssDNA) oligos (IDT, Surrey, UK). TRIB2 oligos were cloned into LentiCRIPSR v2 (AddGene Plasmid #52961) and TRIB1 oligos were cloned into lentiCRISPR v2-Blast (AddGene Plasmid #83840). Lentivirus containing the CRISPR vector was generated by co-transfection of psPAX2 and pVSVG in HEK293T cells using PEI reagent. The CRISPR lentivirus was used to constitutively express sgRNAs and Cas9 under control of human U6 (hU6) and elongation factor 1α (EF-1α) promotors, respectively. Briefly, 2x10^6^ HEK293T cells were seeded in 10cm dishes and incubated for 24hrs.

10µg CRISPR plasmids were diluted in 1ml OPTIMEM media with 7.5µg psPAX2 and 4µg pVSVG using a 3:1 ratio of PEI:DNA. After 20 minutes incubation, the transfection mixture was added to HEK293T cells dropwise. After 24 hours, media was replaced with 6ml fresh medium. At 48hrs and 72hr, the media were filtered, harvested and employed immediately to transduce THP1 cells. CRISPR lentivirus assembled in 48hr and 72hr supernatants was employed on days 1 and 2 respectively, using polybrene and spinoculation, as described for retroviral transduction. At the end of day 2, cells were moved into antibiotic selection medium (2µg/ml puromycin or 24µg/ml blasticidin) and cell counts and viability monitored via trypan blue exclusion assay.

### DNA guides and primer sequences

TRIB1 sgRNA1 fw: 5’-CAC CGG CAG CAG GTA GTC GGC GAT G-3’, TRIB1 sgRNA1 rev: 5’-AAA CCA TCG CCG ACT ACC TGC TGC C-3’, TRIB1 sgRNA2 fw: 5’-CAC CGC GAG CGC GAG CAT GTG TCC C-3’, TRIB1 sgRNA2 rev: 5’-AAA CGG GAC ACA TGC TCG CGC TCG C-3’, TRIB1 sgRNA3 fw: 5’-CAC CGT TTG GCC GGG ACG CCT CGG G-3’, TRIB1 sgRNA3 rev: 5’-AAA CCC CGA GGC GTC CCG GCC AAA C-3’, TRIB2 sgRNA1 fw: 5’-CAC CGG TTG TCG TCT ATA AGG TCC G-3’, TRIB2 sgRNA1 rev: 5’-AAA CCG GAC CTT ATA GAC GAC AAC C-3’, TRIB2 sgRNA2 fw: 5’-CAC CGG AGA TCG CGG AAC AAA ACC C-3’, TRIB2 sgRNA2 rev: 5’-AAA CGG GTT TTG TTC CGC GAT CTC C-3’, TRIB2 sgRNA3 fw: 5’-CAC CGC ATA TCT CGC TAT TGT GAT G-3’, TRIB2 sgRNA3 rev: 5’-AAA CCA TCA CAA TAG CGA GAT ATG C-3’, NTC fw: 5’-CAC CGA GCT CGC CAT GTC GGT TCT C-3’, NTC rev: 5’-AAA CGA GAA CCG ACA TGG CGA GCT C-3’ qPCR: TRIB2 fw primer: 5’-AGC CAG ACT GTT CTA CCA GA-3’, TRIB2 rev primer: 5’-GGC GTC TTC CAG GCT TTC CA-3’

### In vivo engraftment study

To compare engraftment potential *in vivo*, 3x10^6^ retrovirally modified THP1 cells were transplanted via intravenous injection into NOD SCID Gamma (NSG) mice (3 mice per arm of the study). Mice were culled humanely at 27 days post-transplantation. Organs/samples were harvested for analysis by flow cytometry included lymph, spleen, bone marrow and peripheral blood. Samples for flow cytometry were stained with DAPI and anti-human CD45+ antibody (BD Pharmingen™ APC-Cy™7 Mouse Anti-Human CD45) and engraftment was reported as the percentage of double positive (GFP/CD45) cells.

### Quantifications and Statistical Analyses

For quantification of immunoblots, ImageJ software was employed. For quantification of TRIB1 and TRIB2 mRNA levels in cell lines, the DEPMAP portal was used to obtain mRNA transcript expression levels (data version 25Q2). GraphPad Prism software was used for all statistical analyses and for data visualisation and graphical presentation. For comparison between 2 groups, a two-tailed t-test was performed to determine data significance. When comparing 1 variable across multiple groups, a one-way ANOVA or mixed effects model with Holm-Sidak multiple comparison test was used in absence or presence of missing values, respectively. When more than 2 groups with two variables were compared, a two-way ANOVA with Tukey’s multiple comparisons test for significance was used. Significance was attained with reported p-values of <0.05 (*), <0.01 (**), <0.001 (***) and <0.0001 (****).

## Results

### Pharmacological degradation of TRIB2 induces cell death in AML models

The multi-ERBB inhibitor Afatinib was previously identified as a cellular TRIB2 destabiliser after discovery in a high-throughput compound screen (Foulkes *et al,* 2018). To test the potential relevance of pharmacological TRIB2 modulation in AML function and survival we investigated responses to Afatinib amongst a cohort of genetically heterogenous AML cell lines. Following 10µM Afatinib treatment, TRIB2 protein was rapidly degraded in AML cells within 2 hours (**Figure 1A**). Afatinib treatment also led to a dose-dependent increase in cell death after 24 hours as measured by cell viability in all AML lines tested (**Figure 1B**). Interestingly, AML cell lines expressed TRIB2 mRNA at variable levels, but which was inversely correlated with TRIB1 mRNA. However, no association was observed between TRIB1/2 mRNA levels and the degree of cell death induced by Afatinib (**Figure 1B-D**). To evaluate AML cell killing through ‘on-target’ (ERBB receptor) inhibition, we assessed EGFR stimulation with and without Afatinib treatment. We found little phosphorylation of the EGFR downstream target ERK in response to EGF ligand stimulation in AML cell lines U937 and THP1. Furthermore, Afatinib treatment was unable to significantly reduce phospho-ERK levels. These data suggest that ERBB pathway activity is low in AML cells, and that cell death response to Afatinib treatment may be independent of ERBB inhibition (**Supplementary Figure 1**). To determine whether Afatinib-induced cell death instead results from TRIB2 degradation, we introduced the resistance-conferring C96/104S TRIB2 mutant into AML cells. This locus, corresponding to a Cys-rich region in the atypical TRIB2 α-C helix (**Figure 1E**) was previously established to represent the covalent binding interface of Afatinib in TRIB2, preventing drug-mediated TRIB2 degradation in HeLa cells (13). Comparison of retrovirally transduced AML U937 cells expressing wild type (WT) or C96/104S TRIB2 confirmed that the presence of TRIB2 C96/104S significantly reduced cell death induced by Afatinib, and demonstrated a survival advantage after 24- and 48-hour Afatinib treatment in comparison to WT (**Figure 1F-H**). These findings demonstrate that Afatinib-induced degradation of TRIB2 contributes, at least in part, to the mechanistic basis of AML cell death.

**Figure 1.**
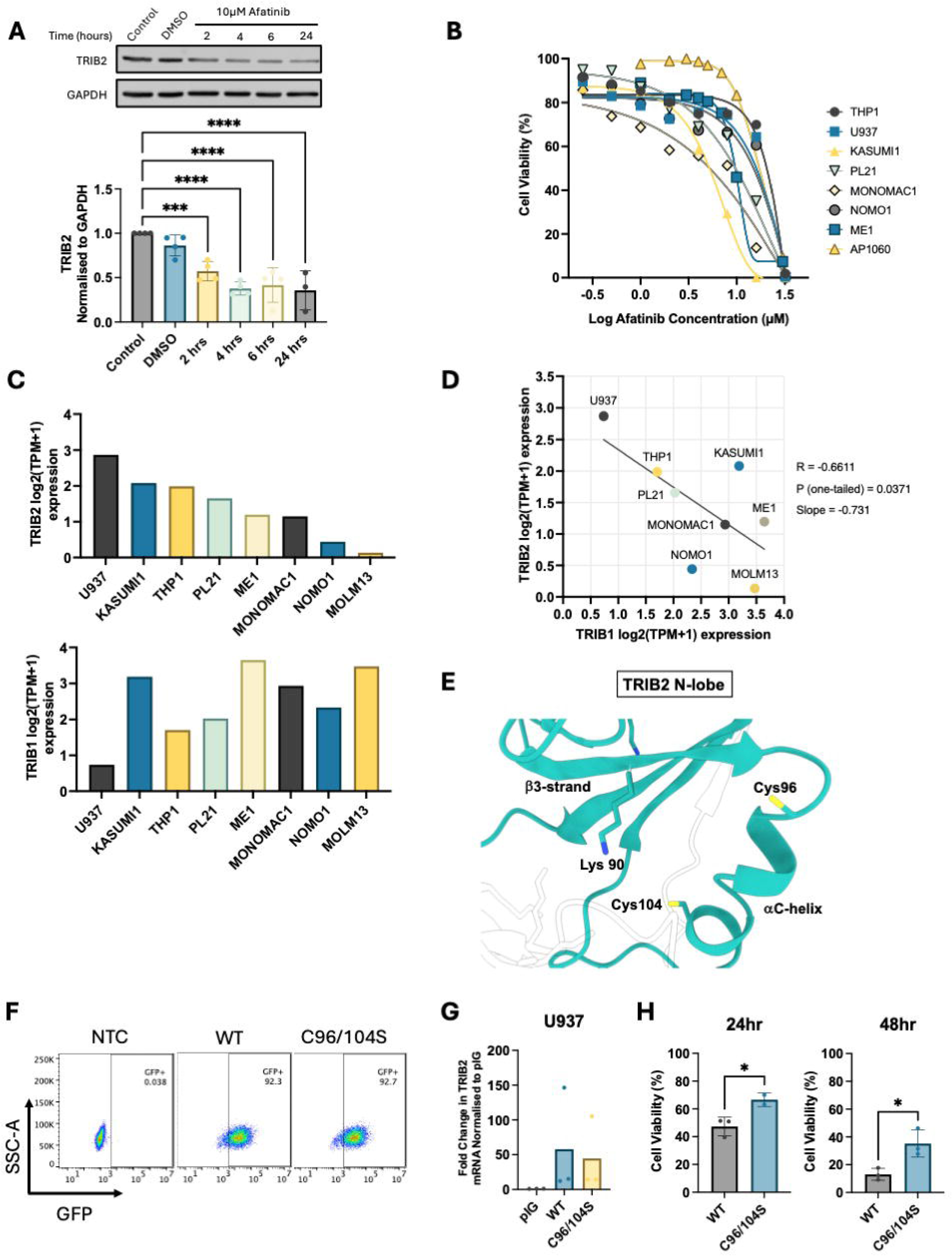
A) Representative western blot (top) and averaged densitometry (bottom) (n=4) of TRIB2 protein levels normalized to GAPDH in AML cell line U937 treated with 10µM Afatinib at indicated timepoints. B) Dose response curve of AML cell lines to Afatinib treatment with cell viability determined via resazurin assay (n=4). C) TRIB2 (top) and TRIB1 (bottom) mRNA transcript expression across AML cell lines as reported by DEPMAP public 25Q2 data version. D) Inverse correlation between TRIB1 and TRIB2 mRNA expression in AML cell lines with Pearson correlation coefficient (R), p-value and slope of the correlation curve given. E) TRIB2 N-lobe with C helix residues Cys-96 and Cys-104 side-chains highlighted in the context of Lys-90, which emerges from the β3 strand in the atypical ATP site. The covalent binding-mode of afatinib has not been established structurally. The predicted alignment error (PAE) for the TRIB2 model is also shown. F) Representative flow cytometry plots of U937 cells transduced with pIG vector containing either WT TRIB2 or C96/104S TRIB2 expression-plasmids showing transduction efficacy of transduced cells compare with negative non-transduced control (NTC) cells. G) qPCR showing fold change in mRNA of TRIB2 in WT and C96/104S TRIB2 transduced U937 cells compare with empty vector control pIG transduced cells (n=3). H) Cell viability of U937 cells transduced with pIG vector containing either WT TRIB2 or C96/104S TRIB2 DNA in response to 16µM Afatinib after 24hr (left) and 48hr (right), assessed by resazurin assay (n=3).

### TRIB2 degradation synergizes with standard AML therapy to induce cell death

The concentration of Afatinib required to elicit maximal responses in AML cells lie within the high micromolar range, making them unlikely to be clinically achievable. We hypothesised that TRIB2 degradation via a multi-ERBB inhibitor could synergise with the standard AML chemotherapy agent cytarabine (AraC) to enhance therapeutic efficacy. We therefore conducted combination treatments across a panel of AML cell lines and used SynergyFinder 2.0 to calculate synergy scores for Afatinib and AraC dose combinations. These data demonstrated enhanced AML cell killing compared to either agent alone, with specific dose combinations resulting in significantly increased levels of cell death. Bliss synergy scores indicated additive effects across all cell lines when averaged over the full dose range. Notably, some dose combinations demonstrated strong synergistic interactions in all but one cell line (**Figure 2A-C and Supplementary Figure 2**). These findings support a potential therapeutic opportunity for small molecule-induced TRIB2 degradation as a complementary therapeutic strategy alongside standard chemotherapy. Whether this is a clinically-viable option with currently approved drugs warrants further investigation.

**Figure 2.**
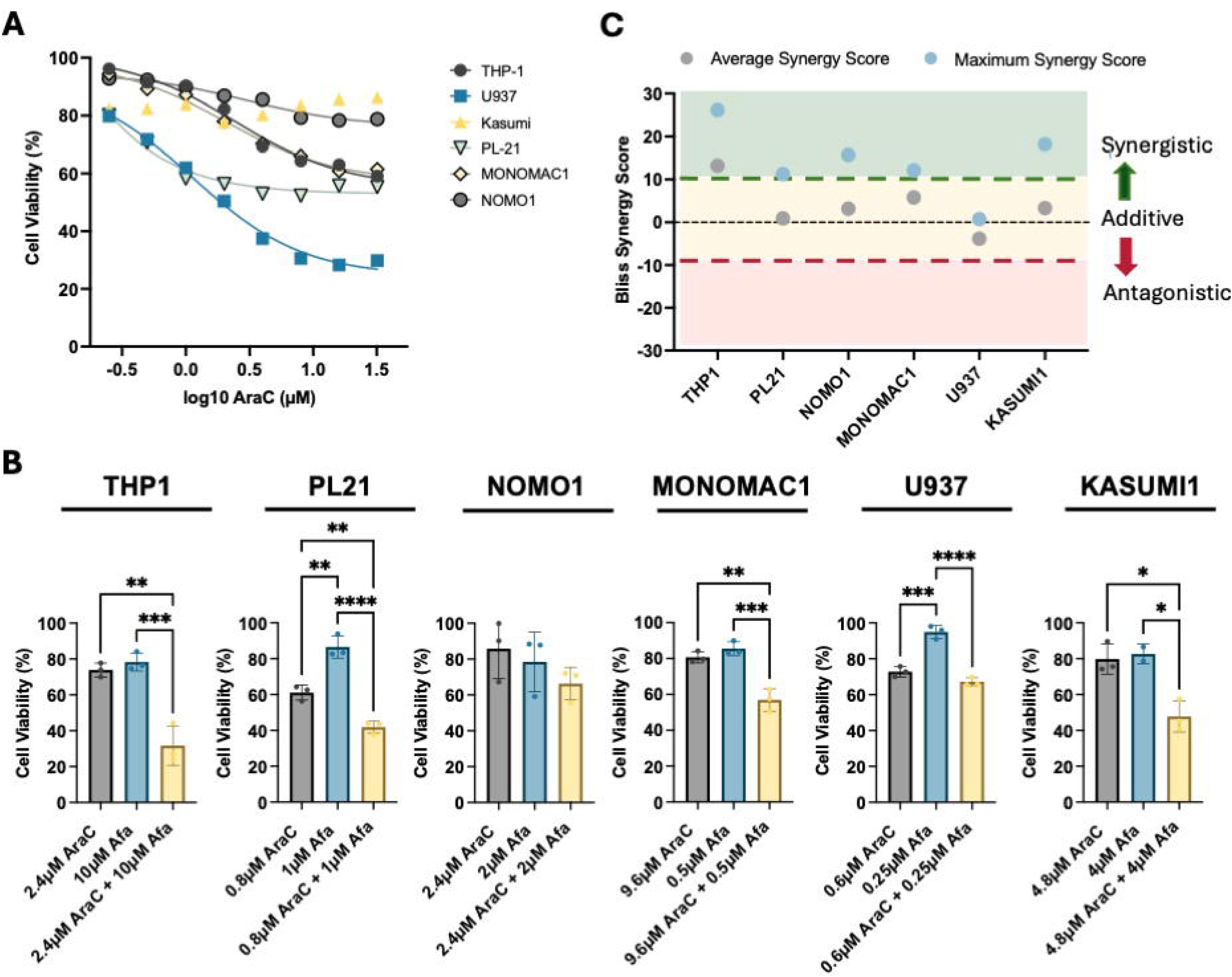
A) Dose response curve of AML cell lines to Cytarabine (AraC) treatment with cell viability determined by resazurin assay (n=4). B) Bar charts showing synergistic dose combinations between AraC and TRIB2-degrading compound (Afatinib) across cell lines, determined via resazurin assay (n=3). C) Summary of average and maximum synergy values of AraC with TRIB2-degrading compound (Afatinib) in each cell line across the drug doses investigated (n=3).

### TRIB2 turnover is mediated by Lys-dependent proteasomal degradation

TRIB2 protein is unstable after isolation *in vitro* (20, 21) and is rapidly turned over in human cells through ubiquitin-dependent mechanisms (22, 23) involving the extended disordered region lying N-terminal to the pseudokinase domain (**Supplementary Figure 3**). Only two sites of TRIB2 ubiquitination are reported in UniProt; Lys-20 and Lys-192, and we previously reported a casual requirement for Lys-177 and Lys-180 residues within the atypical catalytic loop to support C/EBPα degradation by TRIB2 (2). To investigate mechanisms underlying (ubiquitin-mediated) TRIB2 degradation and proteostasis, we performed a mutational screen involving all sixteen TRIB2 Lys (K) residues, which are distributed across the full-length TRIB2 polypeptide (**Figure 3A**), and include Lys-63 in the β-1 strand in the TRIB2 N-lobe, which is predicted to form an interaction with Glu-118 from β-4 in the AlphaFold 3 model **(Figure 3B)**. To identify site-specific requirements for TRIB2 ubiquitination and proteasomal degradation, we first generated a TRIB2 mutant in which all Lys residues were substituted with Arg (R), referred to from here-on as the ‘K*all*R’ mutant (**Figure 3C, left**). To establish site-specific Lys effects in the context of TRIB2 stability, we next individually-changed each Arg residue back to Lys (**Figure 3C, right**). FLAG-tagged TRIB2 expression and ubiquitination assays were performed after immunoprecipitation, with HA-ubiquitin conjugation to FLAG-TRIB2 monitored by immunoblotting. As expected, the K*all*R mutant exhibited the lowest levels of covalent ubiquitination, while the R63K mutant showed the highest levels of ubiquitin conjugation amongst the single-Lys revertants, when compared to wild-type (WT) TRIB2 (calculated as the ratio of total HA-Ub to FLAG-TRIB2 in the immunoprecipitation) (**Figure 3D, E).** These findings suggest that Lys-63, is likely a relevant site for controlling TRIB2 ubiquitination and subsequent proteasomal degradation.

**Figure 3.**
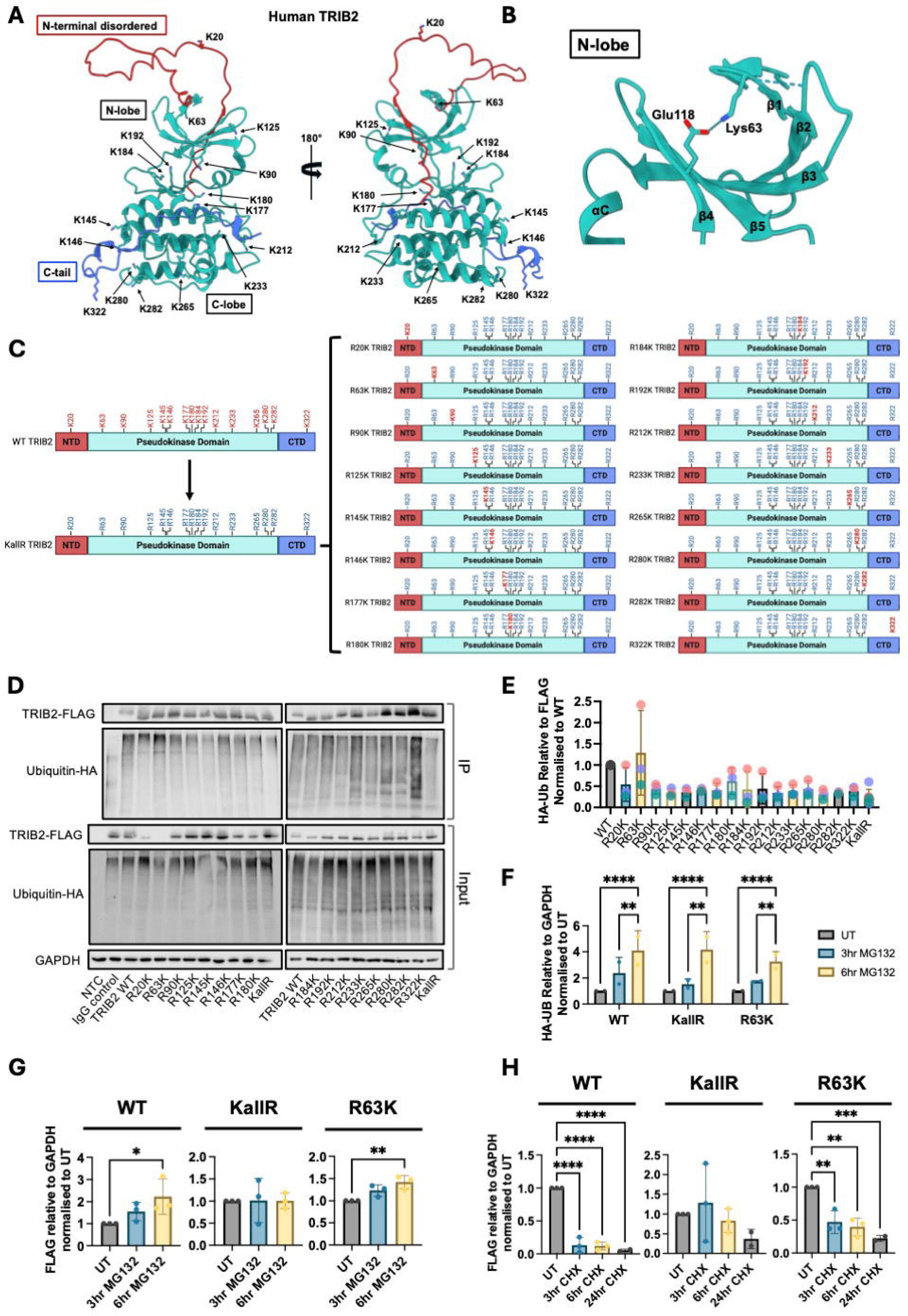
A) AlphaFold 3 full-length human TRIB2 shown in two orientations, rotated 180° along the Y-axis. Lys side-chains are shown as all atom representations. Amino acids composing N-terminal disordered region (1-60) coloured in red, TRIB2 pseudokinase domain (61-308) is coloured teal and C-terminal tail (309-343) is coloured in blue. B) Zoomed-in cartoon of TRIB2 N-lobe, showing Lys-63 in context of Glu-118, with pLDDT scores for the model coloured and graded. Predicted alignment error (PAE) is shown at right. C) Schematic of TRIB2 Lys (K) to Arg (R) substitution library. NTD=N-terminal disordered, CTD=C-tail domain. D) Representative western blots from immunoprecipitated TRIB2 protein after transient expression in HEK293T with HA-tagged ubiquitin. HA-tagged ubiquitin levels were quantified following immunoprecipitation with anti-FLAG antibody to pull down TRIB2 mutants. E) Averaged densitometry (n=3) showing total ubiquitin levels after immunoprecipitation with anti-FLAG antibody to isolate TRIB2 mutants. HA-Ubiquitin protein abundance was normalized to FLAG-TRIB2 and each TRIB2 variant then normalized to WT-TRIB2. Coloured dots indicate matched replicates from each experiment. F) Averaged densitometry (n=2) of HA-ubiquitin in HEK293T cells after transient transfection with relevant TRIB2 plasmid following treatment with 10µM MG132. G) Averaged densitometry (n=3) of FLAG-TRIB2 in HEK293T cells after transient transfection with indicated TRIB2 plasmid following treatment with 10µM MG132. H) Averaged densitometry (n=3) of FLAG-TRIB2 in HEK293T cells after transient transfection with indicated TRIB2 plasmid following treatment with 100µM Cyclohexamide (CHX).

To specifically assess the role of Lys-63 in cellular TRIB2 stability, we compared WT, K*all*R, and R63K TRIB2 protein levels in HEK293T cells treated with the proteasome inhibitor MG132. MG132 increased the total ubiquitin levels detected in all samples (**Figure 3F**). FLAG-tagged WT and R63K TRIB2 revealed similar levels of ubiquitination following MG132 treatment, indicating comparable degradation dynamics. In contrast, MG132 had the least effect on K*all*R TRIB2, consistent with a lack of Lys residues required for ubiquitin-mediated proteasomal targeting (**Figure 3G**), and proving that K*all*R TRIB2 is hyper-stable. Cycloheximide (CHX) ‘pulse-chase’ assays further supported these observations. WT and R63K TRIB2 exhibited protein turnover after 3 hours of CHX treatment, whereas K*all*R TRIB2 remained stable up to 24 hours, indicating resistance to ubiquitin-mediated degradation (**Figure 3H**). Collectively, these data identify Lys-63 as a critical site for TRIB2 ubiquitination and subsequent proteasomal degradation, and confirm that a TRIB2 K*all*R mutant is highly stable due to the absence of Lys residues required for its own turnover.

### Stabilised TRIB2 enhances chemoresistance and engraftment potential in AML cells

To assess the impact of TRIB2 hyperstability on AML progression, we retrovirally transduced THP1 and U937 AML cell lines with GFP-tagged constructs expressing either wild-type (WT) TRIB2, Lys-deficient TRIB2, or degradation-resistant K*all*R TRIB2 mutant. TRIB2 overexpression was confirmed by qPCR and GFP detection via flow cytometry (**Figure 4A-B**). To evaluate chemoresistance, we treated transduced cells with increasing doses of AraC and measured cell viability. Cells expressing K*all*R TRIB2 exhibited enhanced survival compared to those expressing WT TRIB2 in both THP1 and U937 lines (**Figure 4C**), indicating that stable TRIB2 contributes to a chemoresistant AML phenotype. To further investigate the functional consequences of TRIB2 hyper-stabilisation, we performed xenotransplantation experiments using NSG mice injected with WT or K*all*R TRIB2-transduced AML cells. Twenty-seven days post-transplantation, mice were sacrificed, and engraftment was assessed by flow cytometry for GFP and human CD45 in bone marrow and spleen tissues (**Figure 4D**). THP1 cells expressing hyper-stable K*all*R TRIB2 showed increased engraftment in both bone marrow and spleen compared to WT TRIB2-expressing cells (**Figure 4E**). These findings demonstrate that TRIB2 stabilisation (likely via reduced proteasomal degradation) enhances both chemoresistance and engraftment capacity in AML cells. In addition, this provides strong rationale for the development of TRIB2-targeted therapies aimed at overcoming TRIB2 associated drug-resistance to mitigate AML aggressiveness.

**Figure 4.**
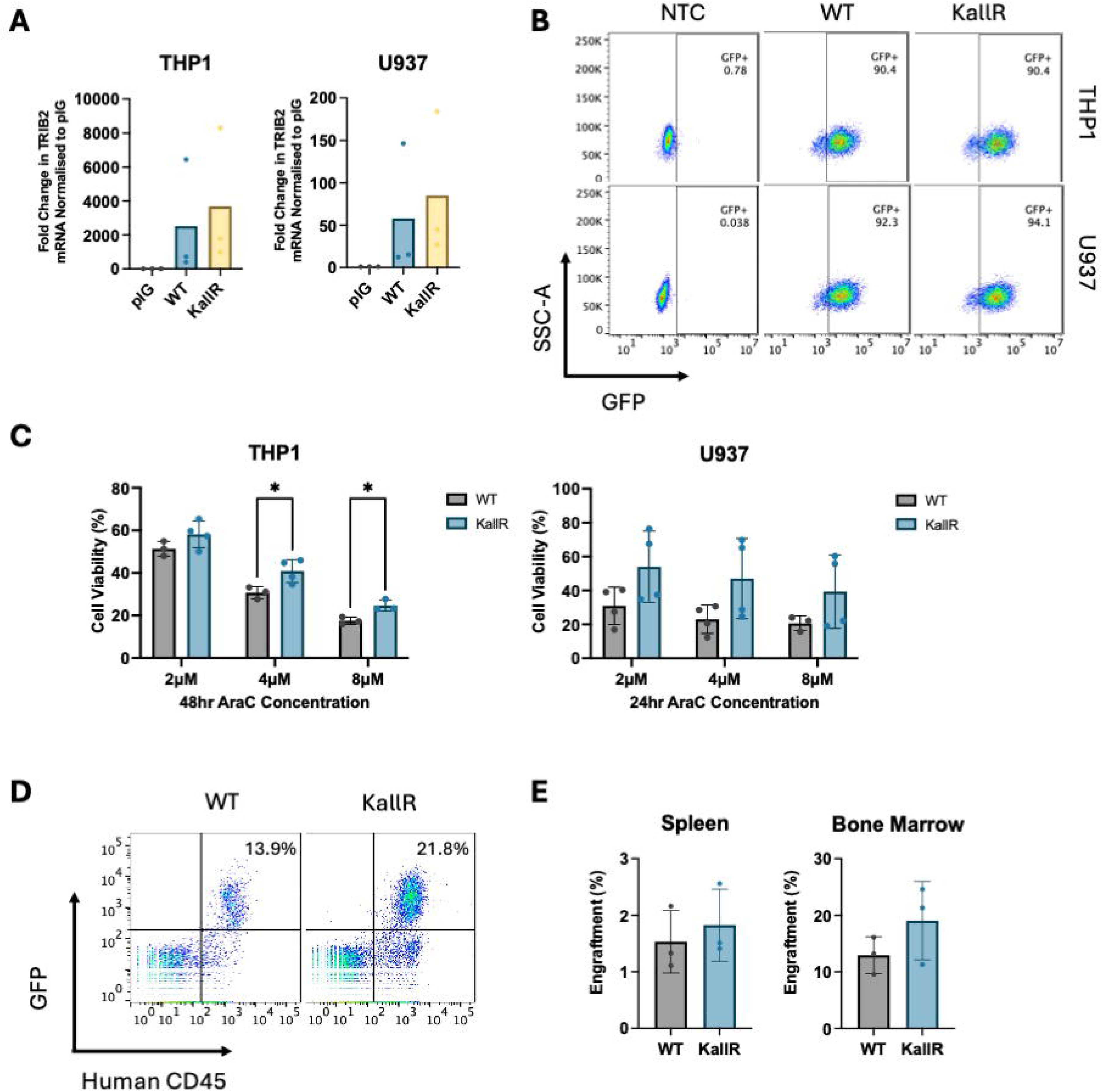
A) qPCR showing fold change in mRNA of TRIB2 in WT and K*all*R TRIB2 transduced THP1 (left) and U937 (right) cells compared with empty vector (control) pIG transduced cells (n=3). B) Representative flow cytometry plots of U937 and THP1 cells transduced with pIG vector containing either WT TRIB2 or K*all*R TRIB2 plasmids showing transduction efficacy of transduced cells compare with negative non-transduced control (NTC) cells. C) Cell viability of THP1 (left) and U937 (right) cells transduced with pIG vector containing either WT TRIB2 or K*all*R TRIB2 plasmids in response to AraC for 48hr and 24hr, respectively. Cell viability determined via resazurin assay (n=4). D) Representative flow cytometry plots showing engraftment levels in the bone marrow of NSG mice engrafted with WT TRIB2 or K*all*R TRIB2 expressing THP1 cells 27 days post-transplantation. Percentage engraftment defined by GFP+humanCD45+ cells. E) Averaged percentage engraftment in the spleen (left) or bone marrow (right) from (D)(n=3).

### Precision Targeting of TRIB2 via PROTACs Induces AML Cell Death

To validate TRIB2 degradation as a viable therapeutic strategy in AML, we first employed a CRISPR-Cas9 gene editing approach to knock out TRIB2 in THP1 cells using three distinct sgRNAs. Following selection of transduced cells, two of the three sgRNAs cell lines significantly impaired cell proliferation and induced cell death, as confirmed by cell counts and trypan blue-based viability assays (**Figure 5A**). While informative, gene editing approaches currently have limited clinical applicability in oncology. Instead, targeted protein degradation (TPD) has emerged as an exciting drug-modality that can effectively target any intracellular protein for degradation. Many TPD modalities, including proteolysis targeting chimeras (PROTACs), function by recruiting an E3 ligase to the protein of interest, promoting its polyubiquitylation and subsequent proteasomal degradation. Given our earlier findings that TRIB2 degradation is Lys-dependent, we explored a TPD strategy using the novel TRIB2-targetting PROTAC compound 5K, which was previously shown to be effective in prostate cancer models (16). We treated a panel of myeloid leukaemia cell lines with increasing doses of the TRIB2-5K PROTAC over a 72-hour time course. All cell lines exhibited dose- and time-dependent cell death following treatment (**Figure 5B-D**). Importantly, TRIB2 protein levels were markedly reduced as early as 6 hours after PROTAC exposure, in advance of cell death onset (**Figure 5E-F**). These results suggest that the TRIB2-5K PROTAC effectively induces AML cell death through targeted degradation of TRIB2, potentially supporting its potential as a novel therapeutic modality in AML.

**Figure 5.**
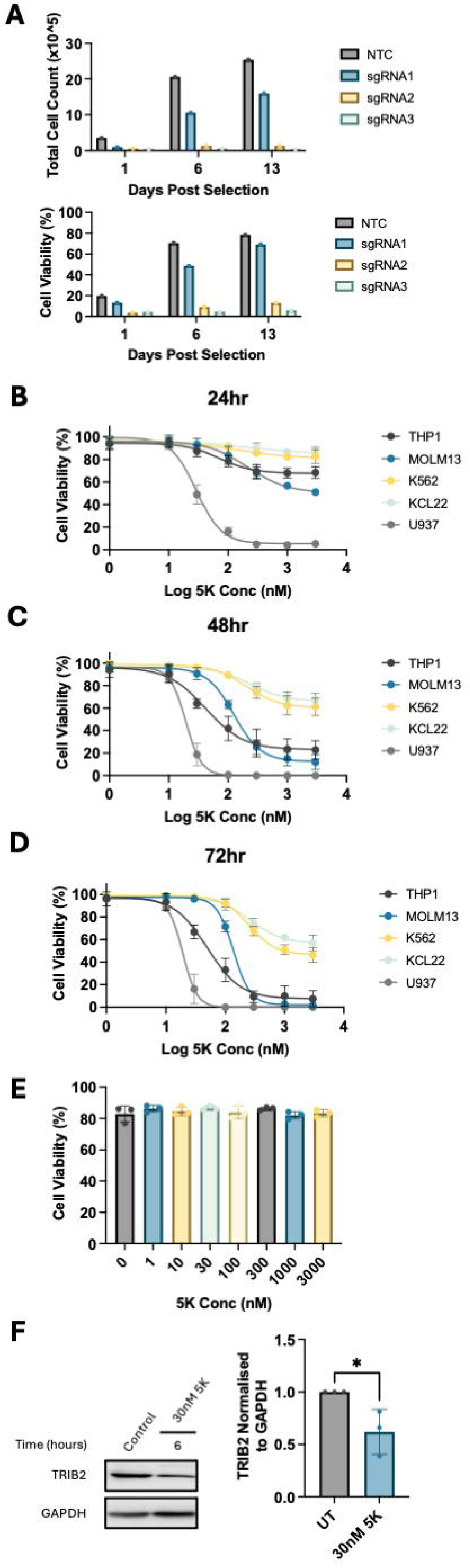
A) Bar graph of cell count (top) and cell viability (bottom) of THP1 cells following TRIB2 CRISPR mediated knock-out with 3 guides (sgRNA1-3) and non-targeting control (NTC). B-D) Dose response to 5K TRIB2 PROTAC treatment over 24hr (B), 48hr n= (C) and 72hr (D) in the indicated leukaemia cell lines. Cell viability determined by resazurin assay (n=4). E) U937 cell line dose response to 5K TRIB2 PROTAC treatment at 6hrs. Cell viability determined by apoptosis assay (n=3). F) Representative western blot and bar graph of TRIB2 expression in U937 cells treated with 30nM 5K at 6hrs.

## Discussion

Despite advances in molecular profiling and targeted therapies, resistance to conventional chemotherapy remains the major driver of poor outcomes in AML. In this study, we demonstrate that targeted degradation of TRIB2 offers a new promising therapeutic strategy. Pharmacological degradation of TRIB2 using Afatinib led to rapid TRIB2 degradation and AML cell death, which was likely independent of ERBB signalling. Mutation of the Afatinib-binding site on TRIB2 (C96/104S) prevented TRIB2 degradation and rescued cell viability, confirming the specificity of the interaction for the phenotype. However, the high micromolar concentrations required for effective TRIB2 degradation by Afatinib likely limit its clinical applicability even though its covalent mechanism of action may help concentrate the drug in cells expressing TRIB2. To address this issue, we explored combination therapy with AraC, revealing synergistic effects that significantly enhanced AML cell killing *in vitro*. These findings support the potential of TRIB2-targeted agents to sensitize AML cells to chemotherapy and overcome resistance. Mechanistically, we identified Lys-63 as a critical TRIB2 site that supports its own ubiquitination and proteasomal degradation. We cannot rule out other Lys residues noted in other literature (UniProt, (2)) such as Lys-20 and Lys-180 having a role. Consistently, mutation of all TRIB2 Lys residues rendered it highly stable, with AML cells expressing hyper-stable TRIB2 exhibiting increased chemoresistance and enhanced engraftment *in vivo*. These results underscore the pathological significance of TRIB2 turnover in AML and further highlight the therapeutic potential of modulating TRIB2 stability *in vivo*.

The general challenge of targeting pseudokinases using conventional ATP-site directed strategies that are readily employed for canonical kinases (24, 25), is ably demonstrated by the dearth of agents that specifically target TRIB2 (26). Repurposing multi-ERBB inhibitors offers one potential translational strategy, with Afatinib demonstrating TRIB2 destabilization in different cell types (5,15) alongside AML cell killing *in vitro* (15). The ability of small molecule kinase binders to degrade their targets is inherent in multiple monovalent protein kinase inhibitors (27, 28). Although only effective at high concentrations, the synergy of the covalent compound Afatinib alongside standard-of-care chemotherapy provides a new proof-of-concept for potential combination approaches. Moreover, the development of the first TRIB2 PROTAC degraders such as compound 5K (16), opens-up new avenues for TRIB2-directed therapeutics. To translate our findings into actionable clinically-viable approaches, we evaluated the TRIB2-specific PROTAC 5K. PROTACs represent a transformative drug modality, which hijack the ubiquitin-proteasome system to eliminate target proteins through proteolysis rather than merely modulate enzymatic function. This approach is particularly valuable for targeting "undruggable" proteins such as pseudokinases and transcription factors (29). Recent advances in PROTAC development have revealed potent activity in haematological malignancies, including AML. Notably, PROTACs targeting oncogenic drivers such as FLT3, BCL-XL, and BTK have entered preclinical and early-phase clinical trials (30). Indeed, FLT3-targeting PROTACs have shown efficacy in FLT3-mutated AML models, offering a strategy to overcome resistance to FLT3 inhibitors (31). Moreover, the BTK degrader NX-2127 is currently in phase 1 trials for relapsed/refractory B-cell malignancies, while other PROTACs such as ARV-110 and ARV-471 have shown promise in solid tumours (30). Our data confirm that 5K induces dose- and time-dependent AML cell death, with TRIB2 degradation detectable within 6 hours, prior to apoptosis onset, suggesting on-target activity. These findings establish TRIB2 as a viable PROTAC target in AML and support further pre-clinical development of TRIB2-directed degraders.

Beyond TRIB2, other members of the Tribbles pseudokinase family, notably TRIB1 and TRIB3, have also emerged as oncogenic drivers in AML and other malignancies. TRIB1 promotes AML progression through degradation of C/EBPα and activation of MAPK/ERK signalling, often cooperating with transcription factors such as HoxA9 (32). On the other hand, TRIB3 stabilizes MYC by shielding it from E3 ligase-mediated degradation, enhancing MYC-MAX transcriptional activity (33). These distinct roles also make TRIB1 and TRIB3 attractive therapeutic targets, although PROTAC strategies for TRIB1 and TRIB3 have not yet been reported. TRIB3-targeting PROTACs have shown initial promise in disrupting the TRIB3-MYC axis, leading to MYC destabilization and tumour suppression (34), whilst TRIB1-targeting PROTACS have not yet been developed. In all these cases, further mechanistic analysis of the targets and specificity of these approaches will be required, with rational PROTAC design, through linker engineering and E3 ligase selection, to maximize therapeutic efficacy.

Despite significant promise, PROTACs face several delivery and pharmacological challenges that may impact treatment efficacy (35). Their high molecular weight and polarity often result in poor oral bioavailability and limited membrane permeability, hindering systemic absorption and tissue penetration, particularly into the bone marrow niche where AML cells reside. Additionally, intrinsic chemical complexity can lead to rapid metabolic clearance and off-target degradation, raising concerns about toxicity and therapeutic windows. These limitations are especially relevant for AML, where effective drug delivery to the haematopoietic compartment is essential for *in vivo* assessment. Further medicinal chemistry optimization of the TRIB2 ‘warhead’, prodrug design, nanoparticle encapsulation, and antibody-PROTAC conjugates could all be explored to improve delivery, stability, and tissue targeting.

In summary, the convergence of mechanistic analysis of TRIB2’s role in AML pathogenesis, validated preclinical therapeutic efficacy, advanced targeting technologies, and the urgent need for novel AML therapeutics creates a unique opportunity to translate TRIB2-targeted approaches into clinical practice. Success in this endeavour could provide new personalised treatment options for AML patients, while establishing a paradigm for targeting other pseudokinases across various cancer types, potentially transforming efforts to treat distinct cancers driven by this "undruggable" class of protein kinase.

## Ethics statement

All animal experiments were carried out in accordance with the Animals Scientific Procedures Act of 1986 and following the University of Glasgow Animal Welfare and Ethical Review Board under Home Office Licence. All experiments were performed under Keeshan’s Procedure Project License (PPL no. PP4496278).

## Data availability statement

All data associated with this study are present in the paper or the Supplementary Materials.

## Acknowledgments

We thank Dr Jake Ritchie and Dr Rachel Craig for their valuable assistance in the laboratory. We are grateful for the support from the facilities, and scientific and technical assistance from flow cytometry staff at Wolfson Wohl Cancer Research Centre and the Biological Services Unit and technical staff, especially Karen Dunn at Cancer Research UK Scotland Institute. For the purpose of open access, the author(s) has applied a Creative Commons Attribution (CC BY) licence to any Author Accepted Manuscript version arising from this submission.

## Funding

This work was supported by Blood Cancer UK project grant 19012 (to K.K. and E.R.), an MRC DiMeN PhD studentship (to J.A.H.), an MRC DTP Precision Medicine PhD studentship (to E.R.), a BBSRC project grant BB/X002780/1 (to P.A.E. and J.A.H.) and BB/T007427/1 (to R.J.C and J.W.) and a Medical Research Council Confidence in Concept grant (to K.K.).

## Author Contributions

E.R., A.N., E.K., J.W., L.L., B.Z., and J.A.H. performed experiments and drafted the figures. K.D., and F. Z., provided essential resources. All authors analysed the data. K.K. and E.R. drafted the manuscript, and all authors approved the final version. P.E., J.A.H., L.R., and R.C. provided expert advice and helped with experimental design. K.K. designed and supervised the project.

## Declaration of interests

P.A.E is co-founder and CSO of Sulantrix Ltd. The remaining authors do not declare any competing interests.

---

**Supplementary Figure 1.**
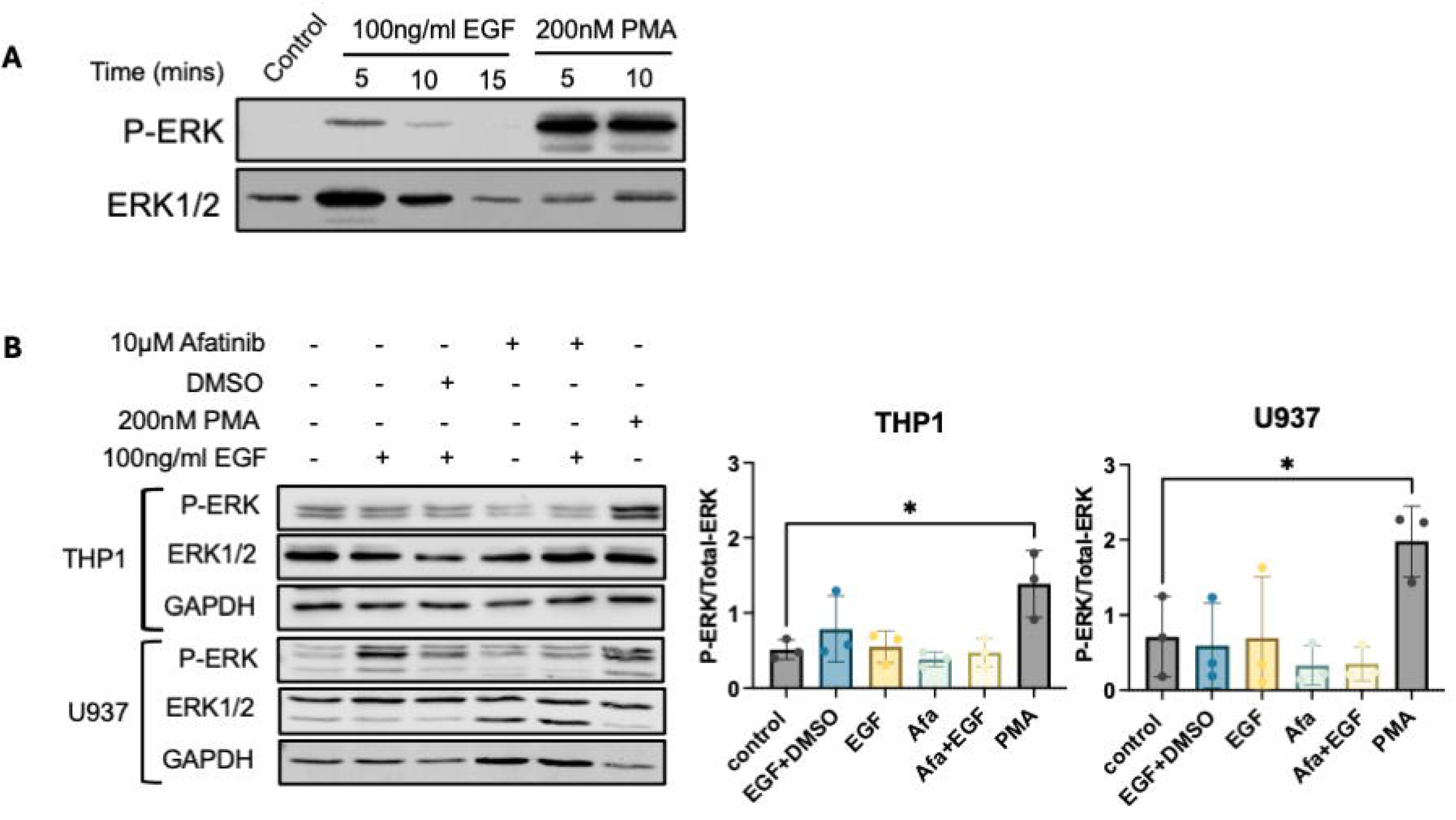
A) Western blot of EGF and PMA stimulated U937 cells showing phospho-ERK relative to total-ERK protein levels. B) Representative western blots and averaged densitometry of EGF, Afatinib and PMA-stimulated U937 and THP1 cells showing phospho-ERK relative to total-ERK protein levels.

**Supplementary Figure 2.**
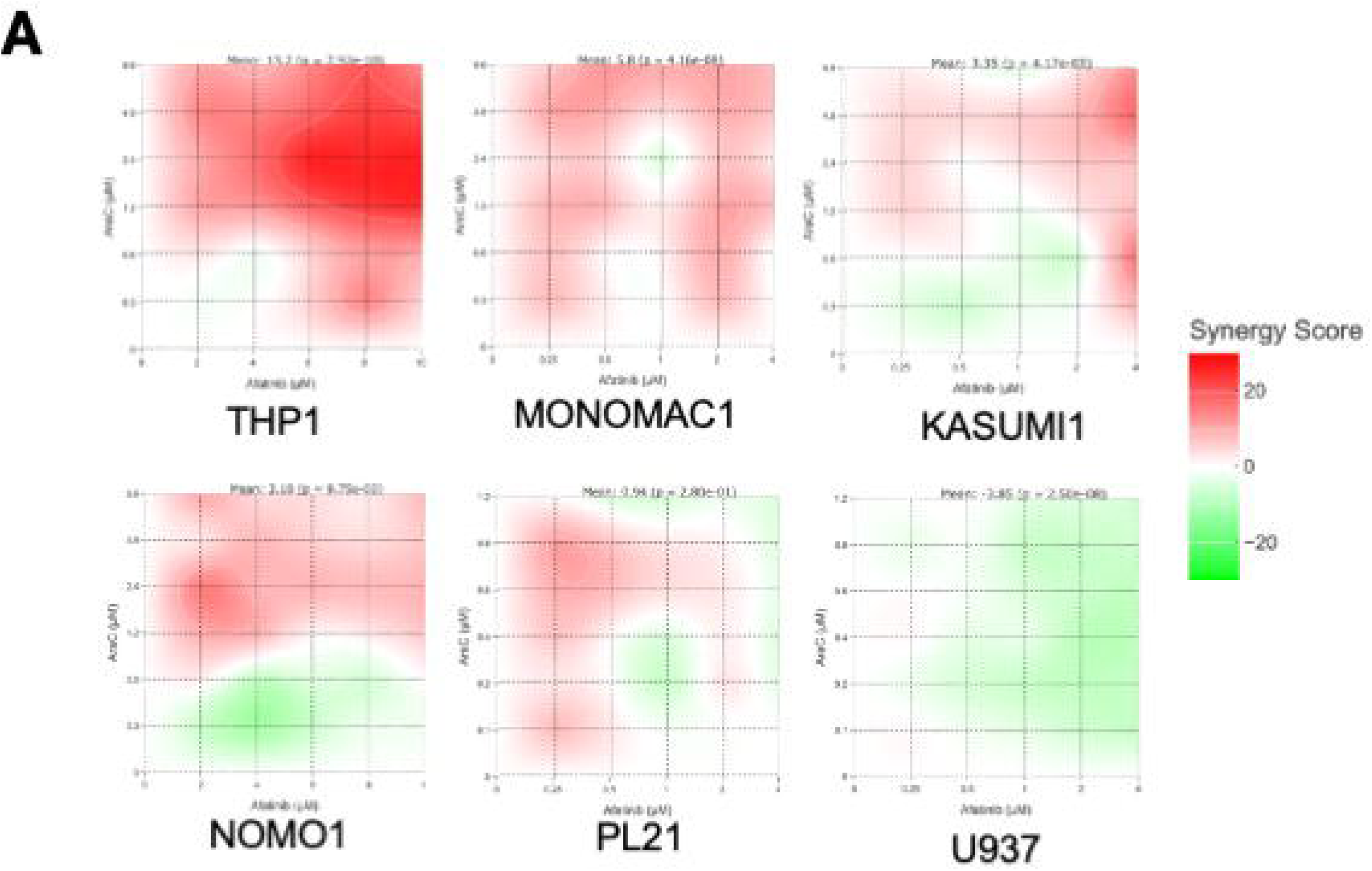
Synergy maps establishing drug synergy between AraC and Afatinib at variable doses in AML cell lines.

**Supplementary Figure 3.**
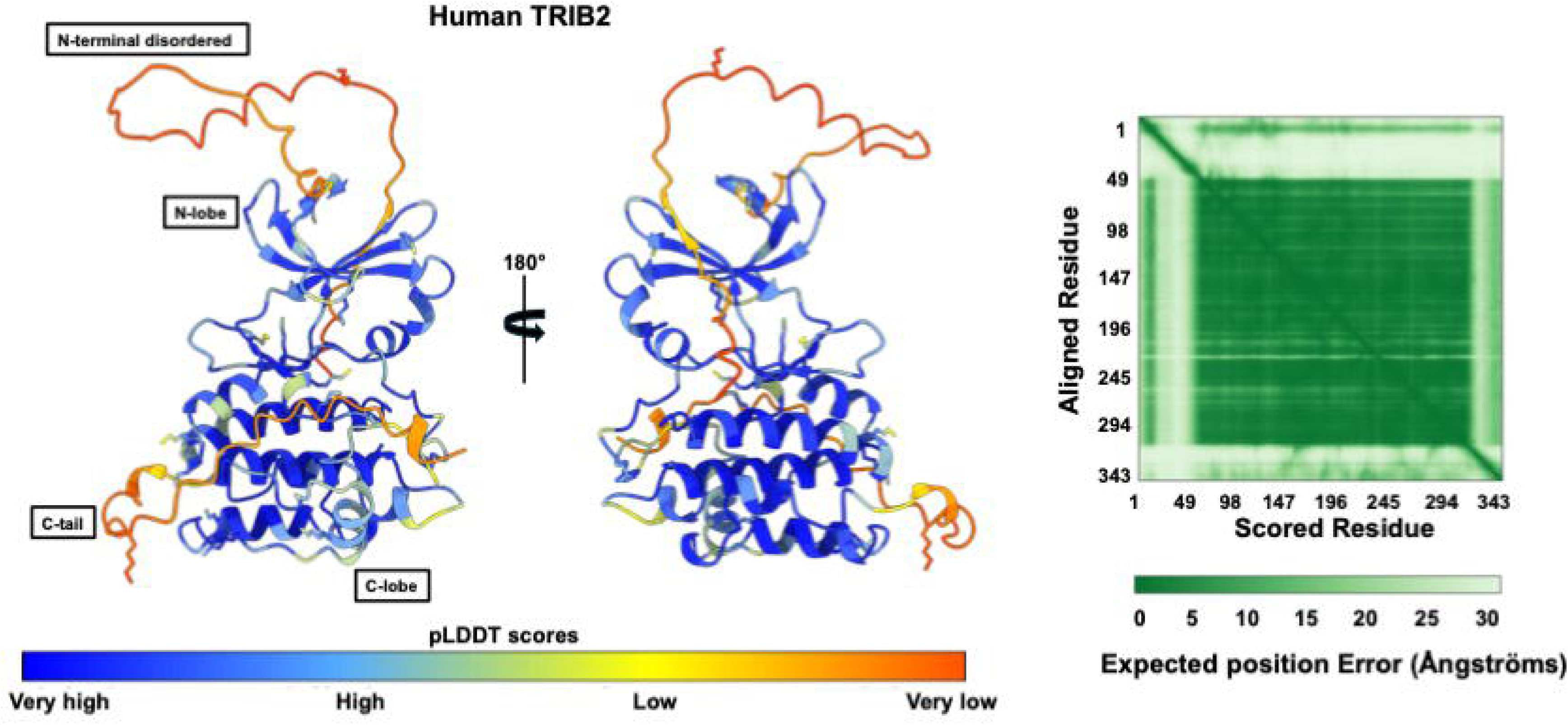
The top-ranked structure of full-length human TRIB2 [1-343] generated using AlphaFold 3 (pTM = 0.77) and visualized in UCSF ChimeraX, coloured by predicted local distance difference test (pLDDT) confidence score. Predicted alignment error (PAE) is shown at right.

